# Protective role of parenthood on age-related brain function in mid- to late-life

**DOI:** 10.1101/2024.05.03.592382

**Authors:** Edwina R. Orchard, Sidhant Chopra, Leon Q.R. Ooi, Pansheng Chen, Lijun An, Sharna D. Jamadar, B.T. Thomas Yeo, Helena J.V. Rutherford, Avram J. Holmes

## Abstract

The experience of parenthood can profoundly alter one’s body, mind, and environment, yet we know little about the long-term associations between parenthood and brain function and aging in adulthood. Here, we investigate the link between number of children parented (parity) and age on brain function in 19,964 females and 17,607 males from the UK Biobank. In both females and males, increased parity was positively associated with functional connectivity, particularly within the somato/motor network. Critically, the spatial topography of parity-linked effects was inversely correlated with the impact of age on functional connectivity across the brain for both females and males, suggesting that a higher number of children is associated with patterns of brain function in the opposite direction to age-related alterations. These results indicate that the changes accompanying parenthood may confer benefits to brain health across the lifespan, highlighting the importance of future work to understand the associated mechanisms.

---

The transition to parenthood is becoming increasingly recognized as a period of considerable neuroplasticity, and a key biosocial developmental life stage for parents themselves^1^. Across pregnancy and the postpartum period, the parental brain undergoes extensive structural and functional plasticity, supporting the requisite behavioral changes associated with caregiving^2–4^. This neuroplasticity is evident in mothers^5,6^ and fathers^7–9^, and is related to both biological and environmental changes, such as hormone levels^10–12^, and the amount of time spent with one’s child^4,9,13^. Although we have gained insight into the initial short-term impacts of parenthood on the brain, the endurance of associated changes in brain function across the lifespan, and their interactions with the process of normative aging remain unexplored.

Decades of research charting age-related brain changes across the lifespan have provided great strides in mapping the normative trajectories of structural^14^ and functional^15^ neural reorganization from early development through late life. Aging in adulthood is associated with a progressive series of functional alterations across sensorimotor and ‘higher-order’ cognitive systems. Typically, connectivity *within* networks, such as the default, salience/ventral attention, and somato/motor networks, decrease with age^15,16^, while connectivity *between* networks increases with age^17,18^. Recent evidence suggests that these network trajectories show different inflection points; for example, compared to higher-order networks, sensorimotor networks reach maturity earlier in life and show a steeper decline in function after the fourth decade of life^14,15^. Although age-related alterations in brain function are broadly consistent across the population, and between males and females^19^, differences in lifelong exposure to biological and environmental factors can impact the timing and rate of these changes, altering aging trajectories. Identifying relevant factors, as well as periods of increased neuroplasticity in adulthood, can highlight windows of vulnerability and opportunity for intervention, as well as increasing our understanding of the normative aging process.

One such sensitive period and relevant factor currently missing from our understanding of normative aging is the impact of reproductive and caregiving experiences on the brain^1^. Importantly, parenthood is an optional life stage that, due to choice, biology, and circumstance, does not occur for all people. Furthermore, the number of children a person has, and their timings, shares complex associations with both biology and sociodemographic factors, including cultural norms and access to contraception^20–24^. Despite these complexities, the most common form of older person is a person who at some stage of their lives became a parent, and as such, our understanding of normative aging is almost entirely based on samples where elderly parents comprise the vast majority. Therefore, a lack of understanding of how parenthood interacts with the aging process also has consequences for our understanding of the life-long health and wellbeing of all humanity, regardless of parenthood status, especially given the rising prevalence of intentions to remain childless in many high-income nations^22,25^.

Theoretical models of cognitive aging and empirical studies of brain function support the hypothesis that parenthood increases environmental novelty and complexity^1,26,27^, increasing cognitive reserve in later life^1^. Recent large-scale population neuroscience studies have demonstrated a consistent picture of the long-term structural brain adaptations related to parenthood. These studies show an association between the number of children a person has parented (parity) and ‘younger-looking’ brain structure^27–33^ and function^26^, in line with earlier work in rodents^34–36^ and non-human primates^37^, suggesting a potentially neuroprotective effect of parity on the human maternal brain, worthy of further study.

Studies of human parenthood often exclude fathers, focusing entirely on gestational mothers. Since males do not experience the same degree of physical and hormonal changes that gestational mothers do in pregnancy, birth, and lactation, examining whether females and males show similar associations between parenthood and age-related brain function can inform our understanding of the biological or environmental mechanisms underlying these effects. For instance, similarity between males and females may suggest that lifestyle factors and environmental changes, experienced by mothers and fathers alike, may play an important role in the association between parity and brain function and implicate the long-term impact of caregiving for parents of all genders who do not experience pregnancy.

Here, we use data from the UK Biobank^38^ – the largest population-based neuroimaging study to date – to comprehensively examine brain function related to parity and aging in both mothers and fathers. We find a widespread pattern of functional alterations, with higher parity associated with increased functional connectivity across the somato/motor network and lower connectivity within cortico-subcortical systems in both males and females, an effect that is highly consistent between sexes. Critically, we find that for both females and males, patterns of parenthood-related brain function are in the opposite direction to those associated with aging, which showed lower functional connectivity across the somato/motor network and higher connectivity within cortico-subcortical systems with age. Our findings closely align with past human^26–33^ and animal^34–37^ work suggesting long-term neuroprotection related to parenthood throughout the lifespan.

## Results

This study used data from 19,964 females and 17,604 males, as part of the UK Biobank^38^ January 2020 release of individuals with complete and useable structural and resting-state functional MRI data, as well as complete information on parity (number of children), age, and sociodemographic measures linked to parity including education and Townsend Deprivation Index. Sample characteristics are provided in Table 1. Our analyses were approved by the Yale University institutional Review Board, and the UK Biobank data was accessed under resource application 25163.

**Table 1.** Sample Characteristics.

| PARITY | 0 |  | 1 |  | 2 |  | 3 |  | 4 |  | 5 |  | 6 |  | TOTAL |  |
| --- | --- | --- | --- | --- | --- | --- | --- | --- | --- | --- | --- | --- | --- | --- | --- | --- |
| SEX | F | M | F | M | F | M | F | M | F | M | F | M | F | M | F | M |
| N | 4482 | 3716 | 2538 | 2155 | 8697 | 7798 | 3343 | 2962 | 734 | 780 | 129 | 151 | 41 | 45 | 19,964 | 17,607 |
| AGE | 52.24<br>(7.30) | 52.96<br>(7.44) | 52.94<br>(7.22) | 54.31<br>(7.77) | 54.6<br>(7.23) | 56.2<br>(7.46) | 55.29<br>(7.2) | 56.57<br>(7.35) | 56.02<br>(7.40) | 57.11<br>(7.39) | 56.1<br>(6.92) | 56.89<br>(7.47) | 56.88<br>(8.74) | 56<br>(7.59) | 54.04<br>(7.35) | 55.39<br>(7.61) |
| TOWNSEND | -1.19<br>(3.01) | -1.11<br>(3.04) | -1.6<br>(2.87) | -1.84<br>(2.77) | -2.24<br>(2.50) | -2.39<br>(2.43) | -2.03<br>(2.55) | -2.19<br>(2.55) | -1.71<br>(2.75) | -1.93<br>(2.58) | -1.44<br>(2.85) | -1.6<br>(2.92) | -1.66<br>(2.91) | -1.23<br>(3.35) | -1.86<br>(2.73) | -1.99<br>(2.69) |
| EDUCATION |  |  |  |  |  |  |  |  |  |  |  |  |  |  |  |  |
| COLLEGE OR UNIVERSITY DEGREE | 55.87 | 54.04 | 45.27 | 49.28 | 45.66 | 51.55 | 47.8 | 54.02 | 44.69 | 56.28 | 41.86 | 47.02 | 39.02 | 51.11 | 48.19 | 52.38 |
| A-LEVELS/AS LEVELS OR EQUIVALENT | 14.90 | 14.96 | 15.48 | 11.88 | 15.25 | 11.99 | 15.5 | 11.58 | 13.35 | 12.44 | 14.73 | 17.22 | 14.63 | 8.89 | 15.17 | 12.59 |
| O- LEVELS/GCSES OR EQUIVALENT | 18.63 | 17.06 | 24.31 | 20.14 | 24.31 | 18.63 | 22.41 | 17.79 | 23.02 | 15.64 | 25.58 | 16.56 | 26.83 | 22.22 | 22.68 | 18.20 |
| CSES OR EQUIVALENT | 3.48 | 4.20 | 5.36 | 5.71 | 4.74 | 4.15 | 4.01 | 3.92 | 4.09 | 3.08 | 3.88 | 5.96 | 4.88 | 8.89 | 4.38 | 4.29 |
| NVQ OR HND OR HNC OR EQUIVALENT | 2.43 | 6.81 | 4.26 | 9.05 | 3.67 | 9.10 | 3.35 | 7.43 | 5.04 | 7.44 | 2.33 | 9.93 | 4.88 | 4.44 | 3.46 | 8.25 |
| OTHER PROFESSIONAL QUALIFICATIONS | 4.69 | 2.93 | 5.32 | 3.94 | 6.38 | 4.57 | 6.94 | 5.27 | 9.81 | 5.13 | 11.63 | 3.31 | 9.76 | 4.44 | 6.13 | 4.28 |
| ETHNICITY (%) |  |  |  |  |  |  |  |  |  |  |  |  |  |  |  |  |
| AFRICAN | .13 | .11 | .16 | .46 | .22 | .17 | .51 | .57 | .54 | 1.54 | 0 | 0 | 0 | 0 | .25 | .32 |
| ANY OTHER ASIAN BACKGROUND | .16 | .30 | .12 | .09 | .08 | .23 | .12 | .27 | 0 | .26 | 0 | .66 | 0 | 0 | .11 | .24 |
| ANY OTHER BLACK BACKGROUND | 0 | 0 | 0 | 0 | 0 | 0 | 0 | 0 | 0 | .13 | 0 | .66 | 0 | 0 | 0 | .01 |
| ANY OTHER MIXED BACKGROUND | .31 | .19 | .20 | .19 | .15 | .08 | .09 | .03 | 0 | 0 | 0 | 0 | 0 | 0 | .18 | .10 |
| ANY OTHER WHITE BACKGROUND | 5.24 | 3.88 | 4.10 | 2.83 | 3.36 | 1.82 | 3.26 | 2.3 | 3.95 | 2.69 | 3.88 | 1.99 | 9.76 | 2.22 | 39.00 | 2.50 |
| BANGLADESHI | .04 | 0 | 0 | .05 | 0 | 0 | 0 | .03 | 0 | 0 | 0 | 0 | 0 | 0 | .01 | .01 |
| BRITISH | 88.11 | 90.23 | 89.36 | 91.14 | 92.31 | 92.97 | 90.7 | 90.68 | 87.60 | 86.54 | 90.7 | 86.75 | 80.49 | 88.89 | 90.51 | 91.44 |
| CARIBBEAN | .60 | .22 | .51 | .46 | .34 | .24 | .36 | .27 | .68 | .26 | 1.55 | .66 | 0 | 0 | .45 | .27 |
| CHINESE | .31 | .24 | .63 | .51 | .32 | .17 | .30 | .17 | .14 | .13 | 0 | 0 | 0 | 0 | .35 | .22 |
| DO NOT KNOW | .02 | 0 | 0 | 0 | .02 | 0 | 0 | 0 | 0 | 0 | 0 | 0 | 0 | 0 | .02 | 0 |
| INDIAN | .33 | .54 | .67 | .88 | .56 | 1.08 | .57 | 1.18 | .54 | 1.15 | 0 | 1.32 | 0 | 0 | .52 | .96 |
| IRISH | 3.30 | 3.20 | 2.56 | 2.04 | 1.66 | 2.21 | 2.66 | 3.00 | 5.18 | 4.10 | 3.1 | 4.64 | 4.88 | 4.44 | 2.45 | 2.64 |
| MIXED | .02 | 0 | 0 | 0 | 0 | 0 | 0 | 0 | 0 | 0 | 0 | 0 | 0 | 0 | .01 | 0 |
| OTHER ETHNIC | .65 | .43 | .91 | .65 | .46 | .35 | .69 | .64 | .68 | .77 | .78 | 1.99 | 4.88 | 2.22 | .62 | .49 |
| PAKISTANI | .09 | .03 | 0 | .09 | .06 | .13 | .12 | .47 | .41 | 1.28 | 0 | 1.32 | 0 | 0 | .08 | .22 |
| PREFER NOT TO ANSWER | .18 | .24 | .20 | .32 | .08 | .33 | .18 | .20 | .27 | 0.90 | 0 | 0 | 0 | 0 | .14 | .31 |
| WHITE | 0 | 0 | .04 | .05 | .05 | .04 | .06 | .07 | 0 | .13 | 0 | 0 | 0 | 2.22 | .04 | .05 |
| WHITE AND ASIAN | .29 | .30 | .08 | .19 | .16 | .09 | .18 | 0 | 0 | .13 | 0 | 0 | 0 | 0 | .18 | .13 |
| WHITE AND BLACK AFRICAN | .07 | .05 | .24 | 0 | .06 | .03 | .03 | .03 | 0 | 0 | 0 | 0 | 0 | 0 | .08 | .03 |
| WHITE AND BLACK CARIBBEAN | .13 | .05 | .24 | .05 | .11 | .08 | .18 | .07 | 0 | 0 | 0 | 0 | 0 | 0 | .14 | .06 |

To enhance comparability and reproducibility, our study used structural and functional MRI data from processing and denoising pipelines designed and carried out by FMRIB, Oxford University, UK^39,40^. Briefly, after the initial minimal processing steps for fMRI data, including gradient distortion and motion correction, intensity normalization, and high-pass temporal filtering, each subject’s data is projected to MNI152 template space^40^. Finally, the data are denoised using FMRIB’s ICA-based X-noiseifier (ICA-FIX)^40,41^. For each individual, the normalized and denoised volumes are used to extract functional time series from 419 parcels, using previously validated 400-region functional cortical^42^ and 19-region subcortical^43^ brain atlases. We then computed pair-wise interregional product-moment correlations, resulting in a 419 × 419 functional connectivity matrix, consisting of 87,571 unique connections per participant.

### Parity is associated with brain function in females and males

To examine the effect of parity on functional connectivity, we computed the Spearman correlation between the number of children parented and functional connectivity at each connection. Given that in adulthood, age, education, and socioeconomic status are associated with both parity^44,45^ and brain function^15,18,46–50^, we adjusted the functional connectivity values for age, education, and Townsend Deprivation Index. This was done separately for females and males. To ensure that our inference was robust to the inclusion of these covariates, we repeated our analyses without covariate adjustment and find consistent results (SFig1). We use the Network Based Statistic (NBS) for family-wise error corrected (FWE) permutation-based inference at the level of connected-components of edges showing a common effect, with significance assessed at *p*_*FWE*_ < .05. To comprehensively map these effects across the 87,571 different connections, we present them across three different scales: (1) the individual connection, or ‘edge’ level (e.g., Fig1A); (2) the network-level (e.g. Fig2A), in which different parcels are aggregated into one of eight canonical brain networks where we show the proportion of implicated connections that fall within or between each brain network, normalized by the total number of possible network connections (see Methods for additional details); and (3) the level of individual functional parcels, to identify specific brain areas housing a high number of implicated connections (e.g., Fig3).

For females, we find a significant and widespread network of 12,740 connections associated with parity, linking 418 parcels (*p*_*FWE*_ < .001; Fig1A-B). The majority of these connections (10,746 connections, 84.3%) show a positive association with parity, in that higher number of children was associated with greater functional connectivity. At a network-level, positive connections were preferentially within the somato/motor network and between the somato/motor, default, visual, and dorsal attention networks (Fig2A-upper triangle), consistent with prior work showing motherhood-related functional alterations in somato/motor regions^5,12,51–55^. Negative connections were concentrated within the subcortex and between the subcortex and the somatomotor network (Fig2A-lower triangle). At the regional-level, positive connections were predominantly located within the bilateral somato/motor cortex, temporal pole, hippocampus, and posterior cingulate cortex (Fig3A), whereas negative connections largely implicate the bilateral thalamus (Fig3C).

**Fig1.**
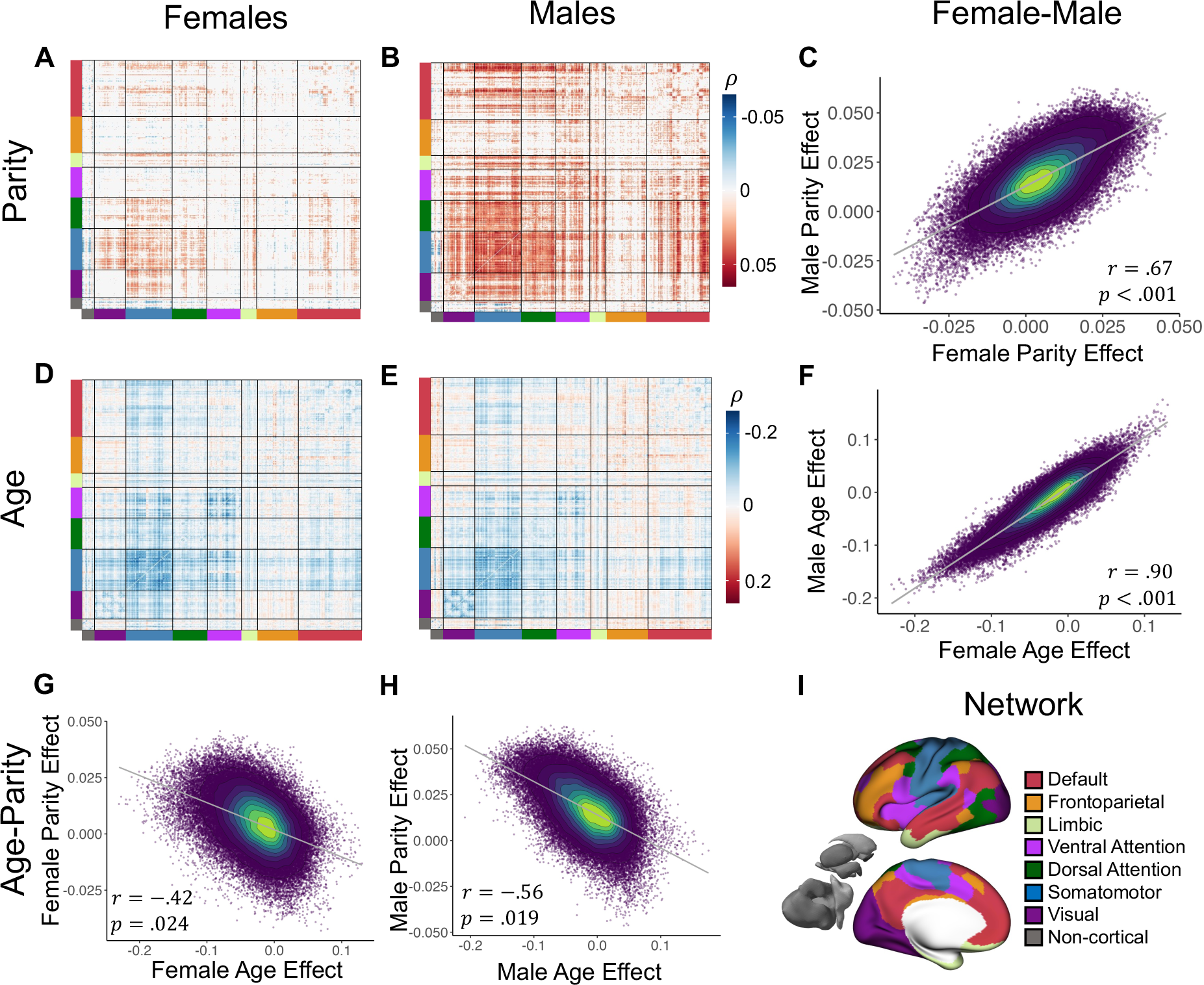
-Effects of parity and age in females and males at the individual connection level. **(A-B, D-E)** 419 × 419 symmetric matrices showing the significant NBS network associated with parity **(A-B)** and age **(D-E)** for females **(A/D)** and males **(B/E)**. Red denotes a positive association, in that higher connectivity was associated with higher parity or age, whereas blue denotes that lower connectivity was associated with higher parity or age. Matrices are ordered by assignment to one of 7 canonical brain networks^56^ and non-cortical regions^57^, represented by corresponding colors at the borders of the matrix, with the network names provided in the lower right legend **(I)**. **(C/F)** Scatter plots of the association between effect sizes across all connections between females and males for parity **(C)** and age **(F)** show high consistency between females and males. **(G-H)** Scatter plots of the association between effect sizes across all connections between the effect of parity and the effect of age on connectivity for females **(G)** and males **(H)** show connectivity effects in the opposite direction for parity and age in both sexes.

**Fig2.**
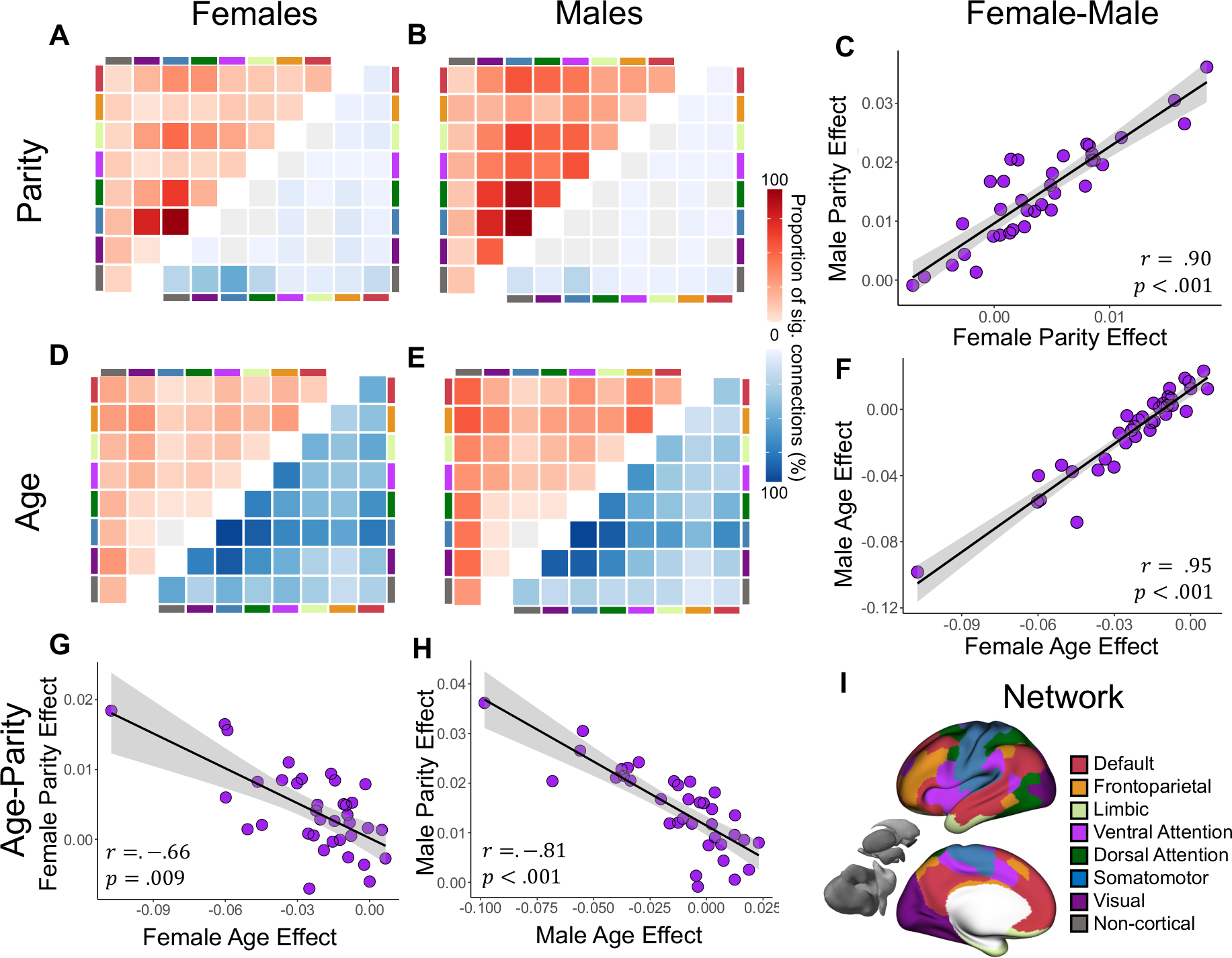
- Effects of parity and age in females and males at the network level. **(A-B, D-E)** Heatmaps of the proposition of connections within the NBS networks which fall within and between each large-scale functional network, normalized for network size (parcel count) associated with parity **(A-B)** and age **(D-E)** for females **(A/D)** and males **(B/E)**. Upper-triangles (red) show network structure of positive connections where higher connectivity was associated higher parity or age, whereas blue (lower triangle) shows negative connections, where lower connectivity was associated higher parity or age. Network assignment is shown by colored bars on the borders, with the network names provided in the bottom right legend. **(C/F)** show the association between effect sizes across all connections between females and males for parity **(C)** and age **(F)**. **(G-H)** Scatter plot of the association between network-level effects between females and males for parity **(C)** and age **(F)**, after averaging the effect size across all connections within and between 8 networks, showing high consistency between females and males. Scatter plot of the association between network-level effects between effect of parity and age for females **(G)** and males **(H)**, after averaging the effect size across all connections within and between 8 networks, showing connectivity effects in the opposite direction for parity and age in both sexes.

For males, we also find a significant widespread network of 36,474 connections associated with parity, linking all 419 regions (*p*_*FWE*_ < .001; Fig1D-E). Similar to females, the majority of these connections (98.4%) show a positive association with parity. At a network-level, positive connections preferentially connected the somato/motor network and between the somato/motor, default, visual, and dorsal attention networks (Fig2B-upper triangle). Negative connections were concentrated within the subcortex and between the subcortex and the somato/motor network (Fig2B-lower triangle). At the regional-level, positive connections predominantly implicated the right insula, bilateral somatomotor cortex, temporal pole, parahippocampal gyrus, and posterior cingulate (Fig3B), whereas negative connections implicated the bilateral caudate, thalamus, and cerebellum (Fig3D).

Given that childlessness is associated with a multitude of genetic^20,21^, socio-demographic^22,23^, and health^24^ factors, we repeated the above analyses after removing individuals reporting 0 children, only including parous females and males, and find highly consistent results at both the individual connection and network levels (see SFig2). This suggests that our results were not driven by the difference between parents and non-parents in our sample.

To examine the consistency of the effect of parity on brain function between females and males, we examined the product-moment correlation between the effect sizes at both the edge level, between 87,571 edges, and at the network level, after averaging the effect size across 36 within- and between-network blocks. We conducted statistical inference using a bootstrapping procedure which preserves the observed association between parity and brain function within each sex^58^. First, both males and females were resampled with replacement 1000 times, and at each bootstrap, the correlation between parity and functional connectivity was computed across all connections for both females and males. Subsequently, we correlated each bootstrapped effect between males and females to build a null distribution of male-female correlations at both the edge- and network level, which was used to obtain a *p*-value.

We find a highly consistent effect of parity on functional connectivity between the sexes, both at the edge (*r* = .67; *p* < .001; Fig1C) and network level (*r* = .90; *p* < .001; Fig2C), suggesting that parenthood may have consistent neurobiological impacts across females and males^15^. This high-level of consistency between the sexes has also previously been demonstrated in age-related structural^14^ and functional^15^ alterations across the life span.

### The effect of parity is in the opposite direction to the effect of aging on brain function

To characterize the effect of age on brain function for females and males, we compute the association between functional connectivity and age at each connection. Give that both education and socioeconomic factors have been consistently shown to impact the association between age and functional connectivity^59,60^, the functional connectivity values were adjusted for education and TDI prior to computing associations with age. We again use the NBS for permutation-based FWE and examine the results across three different scales: 1) individual connections, 2) within and between canonical brain networks, and 3) individual brain regions. We find a strong and widespread effect of age on functional connectivity (*p*_*FWE*_ < .001), for both females (37,076 connections; Fig1D) and males (56,340 connections; Fig1E), linking all 419 regions. Consistent with prior work^15,18^, age was predominantly associated with decreases in functional connectivity, with 32,071 (86.5%) and 38,025 (67.5%) edges showing a negative relationship with age for females and males, respectively. At a network-level, in both females (Fig2D) and males (Fig2E), age-related functional connectivity decreases were concentrated between the somato/motor network and the rest of the brain, whereas increases were concentrated within the subcortex and frontoparietal network. At a regional level, age-related functional connectivity decreases implicated the bilateral somato/motor, temporal and insula cortices, and increases implicated striatal, parietal, and orbitofrontal regions (Fig3E-H). The effect of age on functional connectivity was highly consistent between females and males both at the edge (*r* = .90; *p* < .001; Fig1F) and network levels (*r* = .95; *p* < .001; Fig2F)^30^.

We hypothesized that if parenthood confers a protective effect on age-related decline in brain function, the effect of parity on functional connectivity would be negatively associated with the effect of age on functional connectivity. To test this, for males and females, we examined the product-moment correlation between the effect sizes of parity and age at both the edge level, between 87,571 connections, and at the network level, after averaging the effect size across 36 within- and between-network blocks. We again conducted statistical inference using a bootstrapping procedure which preserves the observed associations between parity and brain function, age and brain function for each sex^58^. During this procedure, for each sex, individuals were resampled with replacement 1000 times, and at each bootstrap, the correlation between parity and functional connectivity, and age and functional connectivity were computed across all connections. Subsequently, we correlated each bootstrapped effect between parity and age to build a null distribution of parity-age correlations at both the edge- and network-level for each sex, which was used to obtain a *p*-value.

Critically, we find that at the level of brain-wide individual connections, the effects of parity on functional connectivity were indeed negatively correlated with the effects of age on functional connectivity both for females (*r* = −.42; *p* = .024; Fig1G), and males (*r* = −.56; *p* = .019; Fig1H), suggesting that higher number of children is associated with patterns of brain function in the opposite direction to age-related brain function in parents of both sexes. When examining this association at a network-level we again find a consistent and strong negative association between the effects of parity and age on functional connectivity in both females (*r* = −.66; *p* = .009; Fig2G) and males (*r* = −.81; *p* < .001; Fig2H).

**Fig3.**
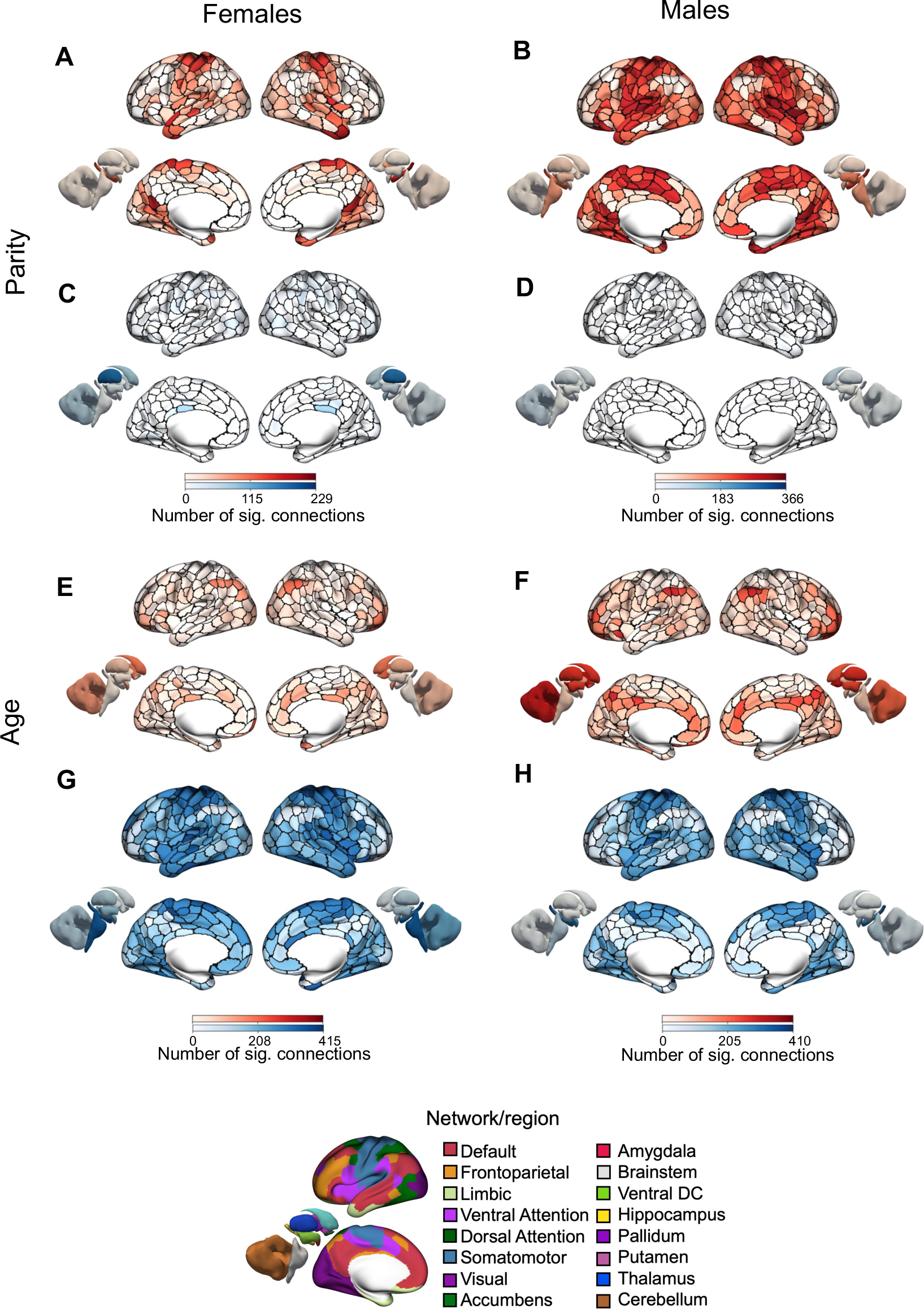
- Brain regions associated with parity and age in females and males. Surface renderings depicting the number of connections (i.e., nodal degree) in the NBS network attached to each brain region for models examining the association between brain function and parity **(A-D)** or age **(E-F)** separately for females (left column) and males (right column). Number of positive connections, where higher connectivity was associated with higher parity or age are shown in red, whereas number of negative connections, where lower connectivity was associated with higher parity or age, are shown in blue.

## Discussion

The transition to parenthood involves rapid, profound, and interacting adaptations that require a flexible restructuring across physical, mental, social, and environmental domains, altering myriad facets of a person’s life. However, the ways in which this sensitive period impacts the brain long-term, and interacts with the aging process, are largely unknown. Here, we investigated the enduring impact of number of children parented (parity) on brain-wide functional connectivity, in a large sample of adults from the UK Biobank^38^. We find a widespread pattern of connections related to parity, that is highly consistent between females and males, suggesting that the impacts of parenthood on human brain function are long-lasting, cumulative, and likely environmentally driven. Critically, the direction of the observed effect of parity on connectivity is in the opposite direction to the effect of age on connectivity, suggesting a potentially protective effect of increased parity on the adult brain, for both females and males. Here, we discuss the potential for these effects to be driven by a combination of 1) direct/caregiving, 2) indirect/environmental, and/or 3) covarying/sociodemographic mechanisms. While the factors underpinning this neuroprotective effect require further study, these results are consistent with the extant human and animal parental brain literatures, which point to structural^27–33^ and functional^26^ neuroprotection in mothers^26–33^ and fathers^30^ with more children.

### Parity is associated with brain function

We find a significant widespread association between parity and functional connectivity in mid- to late-life adults, suggesting that the impact of parenthood on brain function endures across the lifespan. This effect of parity is consistent between females and males, implicating mechanisms of the shared parental environment, rather than solely biological alterations related to pregnancy, birth, and lactation. For both females and males, higher parity was associated with higher functional connectivity, largely concentrated within the somato/motor network, and between the default network and the rest of the brain. For mothers, somato/motor regions are consistently activated in response to infant stimuli^51^, show restructuring of grey^12,52,53^ and white^52^ matter during pregnancy, and grey matter in early motherhood^5,54^, and emerge as functional “hub” regions in first-time pregnant mothers compared to non-mothers^55^, suggesting that somato/motor areas may be impacted by, or important for maternal caregiving. Further, we find that at a regional level, the hippocampus emerges as a highly implicated area, with increased connectivity to the bilateral hippocampi associated with increased parity for both females and males, extending to the bilateral parahippocampal gyri for males. The hippocampus shows considerable structural reorganization across pregnancy^2,12^ and the postpartum period for both mothers^2,54,61^ and fathers^62^, and shows associations with parity for mothers in middle^29^ and late life^27^. For mothers, lower hippocampal volume at four months postpartum is associated with positive mother–child interactions^63^, and connectivity with the parahippocampal gyrus is related to maternal attachment and self-efficacy at one-year postpartum^64^, suggesting structural and functional hippocampal changes have broad implications for caregiving behavior.

While these effects could indicate neural changes resulting from the early stages of parenthood that stabilize and endure throughout the lifespan, they could also indicate networks and regions that are impacted at later stages. As we show an association between parity and brain function, this suggests a cumulative impact of multiparity on brain function, such that having additional children continues to alter brain function in a ‘dose-dependent’ manner. In support of this, we find consistent results when excluding individuals without any children, suggesting that our findings do not simply represent differences between those who are and those who are not parents. Importantly, while we adjusted our models for factors such as education and socioeconomic deprivation, the differences in connectivity associated with parity could be further related to other complex sociodemographic and lifestyle factors that are associated with a greater number of children (discussed below in *Potential Mechanisms*).

Of note, while the observed parity-related functional networks are highly similar between females and males, the associations are generally numerically stronger for males compared to females. This result may reflect biological differences between the mothers and fathers in this sample (e.g., the impact of pregnancy, birth, and lactation) or differences in expectations of caregiving responsibilities between genders^65^. This difference in strength could also reflect more general sex differences in brain function, that are not specifically related to the context of parenthood^66–69^. Taken together, the present study is the largest investigation of parental brain function to date and the first to show differences in parental brain function in males beyond early fatherhood. These results suggest that parenthood is a key biosocial life stage with long-term impacts on brain function for females and males and highlight the importance of further study of the impact of parenthood on the male brain, as well as the impact of caregiving in the absence of pregnancy more generally (i.e., non-gestational parents of all genders).

### Age is associated with brain function

Consistent with a large body of existing aging literature^15,18^, we find that older age was associated with a widespread network connectivity alterations in both females and males^15,18^. While the effect of aging on functional connectivity varies depending on the in-scanner task and methodology used^18^, two of the most consistent findings are that aging is associated with decreasing connectivity, and specifically decreased within-network and increased between-network connectivity^16,18^. The results reported here are consistent with both these canonical findings. Further, we find at the network-level, age-related functional connectivity decreases were largely concentrated between the somato/motor network and the rest of the brain, whereas increases were concentrated within the subcortex and frontoparietal network. Age-related decreases in sensorimotor network connectivity have been consistently reported^70–75^, although not in all studies^76,77^. Moreover, this pattern of dysconnectivity has been described in studies characterizing normative ageing trajectories^15,18^, where the dysconnectivity of the somato/motor network is among the first age-related functional brain changes, consistent with our mid- to late-life sample. Also, in line with previous large-scale studies^15,76^, we find a broad pattern of consistency between sexes in age-related functional changes at the individual connection and network level.

### Parity and age have contrasting effects on brain function

Strikingly, the direction of the observed parity and age effects were negatively correlated, such that the connections that showed higher connectivity with parity, showed lower connectivity with age. This effect is strong at an individual connection-level, and even stronger at a network-level, and exists for both females and males, suggesting a neuroprotective effect of parenthood on brain function in later life. This finding, though striking, is in line with our hypothesis, as well as the current understanding of the enduring impact of parenthood on the parental brain in humans^27–32^ and animals^34–36^. Indeed, a growing literature examining brain structure in later life parenthood demonstrates ‘younger-looking’ brains in adults with more children^27–33^, painting a consistent picture of parity-related protection for the structure of the adult brain. These studies similarly show an association between parity and ‘younger-looking’ grey matter^27–31^ and white matter^32,33^ for mothers, as well as grey matter for fathers^30^. In gestational mothers, similar evidence of this ‘younger-looking’ brain structure has been shown as early as 4-6 weeks following birth^78^. These studies support earlier work in both rodent^34–36^ and non-human primate^37^ models, which show similar benefits to brain anatomy. In the only study of late-life parental brain function^26^, mothers with more children also showed patterns of brain function in the *opposite* direction to three theoretical patterns of age-related decline, again consistent with the interpretation of parity as neuroprotective for human brain function. Taken together, the previous and current results support proposed frameworks for understanding parenthood as a developmental life stage, creating an enriching parental environment and potentially contributing to increased cognitive and neural reserve in middle and later life^1^. The present results bolster our current understanding of the enduring impact of parenthood on the human brain and comprise the first examination of parental brain function in mid- to late-life.

### Potential Mechanisms

The current results suggest the presence of age-related neural benefits with increased parity. However, the associated mechanisms are unknown, and we are left with the question of why this effect might exist. Here, we discuss the potential for these effects to be driven by 1) direct/caregiving, 2) indirect/environmental, and/or 3) covarying/sociodemographic mechanisms.

First, the parental brain may be altered directly via biological changes^10–12^ and/or the behavioral actions of caregiving^4,13^. As mentioned above, many previous studies have also found increased neuroplasticity in somato/motor regions across early parenthood in humans^5,12,51–55^ and animals^35,79,80^. This has been interpreted as resulting from caregiving as a highly sensory, tactile, visual, and auditory process, involving planned and coordinated motor outputs – e.g., cuddling, cradling, and feeding. New parents must become attuned to their child by integrating multiple domains of nonverbal cues, including gaze, facial expression, vocalization, and touch^81^, in order to sensitively respond to their child’s changing needs. This may result in the honing of sensory, visual, tactile, and auditory networks in the parental brain over time^12^. Indeed, work in rodents has shown the fine-tuning of the primary sensory cortex (S1) in lactating dams, where the cortical representation of the nipple-bearing skin increased twofold, compared to non-lactating dams and virgin females^79^, highlighting the sensitization of the sensory circuits caused by the stimulation of nursing. Long-term changes to sensory representations in gestational human mothers have also been self-reported, with up to 40% of an Australian sample reporting experiences of ‘phantom kicks’, where ‘kick-like sensations’ are convincingly felt years after giving birth^82^. Additionally, increased somato/motor network connectivity in males may relate to fathers’ involvement as more stimulatory and intrusive (i.e., the physical manipulation of child’s body/ moving child in space), ‘rough and tumble’ play, and in later years, exploration of the environment and skill learning^83,84^. As this relationship exists for both males and females, it strongly implicates the parental environment, as a shared mechanism for females and males. However, there may also be distinct mechanisms with converging hormonal^10–12^ and/or caregiving^4,9,13^ outcomes similarly impacting increased connectivity in mothers and fathers. Regardless of the mechanism itself (environmental, hormonal, or some mixture of both), a direct impact of caregiving could imply that these effects may also exist for other types of caregivers, with a potential impact for non-gestational parents of all genders, and perhaps even grandparents, childcare workers, and any other person with a strong relationship with, or responsibility for, children.

In addition to direct caregiving mechanisms, these results could also be explained by indirect mechanisms of the parenting environment on lifestyle. For example, increased functional connectivity of the somato/motor network may also reflect increased physical activity or social connectedness and stimulation^85^ in parents with more children (higher parity), as well as additional support provided to parents by their children in later years^86,87^. Increased physical, leisure, and social activity are known to increase functional connectivity in both somato/motor and hippocampal regions, providing a potential mechanism for the present results^88,89^. Furthermore, the increased novelty, complexity, and cognitive challenge inherent in the caregiving environment may provide a form of environmental enrichment for parents, which when sustained across the lifespan, might be beneficial for neural and cognitive resilience in later life^1,34,35,90,91^. Therefore, the observed protective effect might arise from increased neural reserve related to the lifetime cognitive load of the caregiving environment^1^.

In addition to these direct and indirect parenthood mechanisms, the present results could also implicate a number of complex relationships with other relevant variables that covary with parenthood. For example, the number of children a person has parented is influenced by biology (i.e., fertility/virility^20,21^) and sociocultural factors, including their desire to be a parent^92^, cultural norms surrounding birth timing (parental age at each pregnancy) and family size^25^, and access to contraception and reproductive healthcare^93^. Importantly for the study of brain function, lifetime labor force participation, socioeconomic status, and educational attainment all show associations with parity^25,85,93,94^. In our present analyses we adjusted our models for differences in education and socioeconomic deprivation across the study population. Although we acknowledge the inability of covariates to fully capture the nuance and complexity of the human experience, we conducted our primary analyses with and without the inclusion of these covariates and find consistent results. Additionally, the associations between these sociodemographic variables and parity would not necessarily imply the protective impacts that we show here. For example, increased parity is associated with lower socioeconomic status, less labor force participation, and lower educational attainment^93,95^, factors that are not protective for brain aging, suggesting that the protective impact of parenthood on age-related brain function shown here may be over and above these potential associations.

### Limitations and Future Directions

Importantly, just as it is difficult to disentangle the mechanisms by which parenthood exerts an influence on brain function–a direct effect of caregiving, an indirect effect of the parental environment, or at the level of covarying sociocultural relationships–we must be aware of how each of these mechanisms differ between individuals geographically, culturally, and socio-demographically. For example, differences between families, such as the role of extended family and grandparents, multigenerational households, parenting styles, and parental expectations and obligations may all impact both the direct effects of caregiving, as well as any indirect or other covarying effects. This diversity in caregiving experience also exists generationally, for example with recent shifts towards smaller family size, pregnancy postponement (later age at first pregnancy), and increasing numbers of people choosing not to become parents, in many high-income countries^22,93^. With these changing norms in mind, the impact of parenthood on future generations may also change over time. Taken together, it is important to recognize that much of our understanding of long-term changes in the human parental brain has largely relied on data sourced from generally western, educated, industrialized, rich, and democratic (WEIRD) samples^96^. Given the cultural and contextual constraints of the UK Biobank data used here, it is possible that different associations between parity, age, and brain function may be found using alternate and more diverse samples, especially given known associations between the parental brain and differences in stress exposure^97^, socioeconomic disadvantage^98^, psychopathology^99^, and substance use^100^.

Finally, our study defines parity based on biological terms: ‘*number of live births*’ for females and *’number of children fathered’* for males. This narrow definition does not encompass the complexities of parenting roles, diverse family structures, and different pathways to becoming a parent. The study also lacks data on relevant caregiving factors, including attachment and parental involvement. Future research should collect rich information on caregiving, as well as data on non-gestational parents of all genders and people who have experienced pregnancy without caregiving. These data, combined with longitudinal designs, will help disentangle possible mechanisms of parenthood-related neural adaptations.

## Conclusion

In a large population-based sample, the present analyses reveal a widespread association between parity (number of children) and the intrinsic functional architecture of the human brain that is consistent across females and males. Critically, the functional correlates of parity are in the opposite direction to the effect of age on brain function. These findings align with human and animal research in suggesting that the complex biological and environmental effects of parenthood may confer protective advantages to brain function across the lifespan. These results provide a crucial step to understanding the long-term impact of parenthood, and call for future large-scale, prospective, diverse, and longitudinal research to explore and further disentangle the mechanisms of these effects.

## Methods

### Participants

This study used data from the UK Biobank^38^, which is a population epidemiology study of 500,000 adults aged 40–69 years, registered with the National Health Service and recruited between 2006 and 2010. A subset of 100,000 participants was recruited for multimodal imaging, including brain structural MRI and resting state fMRI (rs-fMRI). Here, we used the January 2020 release of 37,848 participants with complete and useable structural MRI and rs-fMRI. Of these individuals, we included participants with complete information on age, sex, parity (number of children), education, and socioeconomic deprivation (see Table 1). Our analyses were approved by the Yale University institutional Review Board and the UK Biobank data was accessed under resource application 25163.

### Parenting and demographic data

These data were collected when participants attended the assessment center for an MRI scan and were downloaded from the UK Biobank Application Management System. Sex in the UK Biobank data was determined based on participant responses to a questionnaire offering two options: female or male. This measure primarily reflects biological sex rather than gender as a social construct, which was not explicitly captured in the UK Biobank. Consequently, in our analysis, we use the terms ‘female’ to refer to individuals who self-identified as female, and ‘male’ for those who self-reported as male.

Parity was determined as the self-reported response to the question *‘How many live births did you have?’* for females and ‘*How many children have you fathered?*’ for males. In both cases, included answers ranged from 0 to 6, with values above 6 excluded from analysis, consistent with other population-based parity studies^26,27^. We repeated our primary analysis after excluding individuals with 0 children to ensure that our findings were not driven by differences between those with and without children (see Supplement).

Age was calculated as age at attendance to the assessment center. Level of education including the attainment of a tertiary degree was determined at first assessment. The Townsend Deprivation Index (TDI) is a tool used to assess socioeconomic deprivation, deriving its values from national census data. It assigns a deprivation score to each participant or geographical area, based on four key indicators: unemployment rate, non-car ownership, non-home ownership, and household overcrowding.

### Brain MRI acquisition and processing

We used the processed volumetric resting-state functional MRI (rs-fMRI) data from the first imaging visit^40^. Briefly, each fMRI dataset was spatially normalized to MNI152 2-mm template space and FMRIB’s ICA-based classifier (FSL-FIX; ^41^) was trained on holdout set of participants and applied to the remaining participants to denoise the data. A detailed outline of the processing, denoising and quality control of these data has been previously reported ^40^. Using previously validated cortical^42^ and subcortical^43^ atlases for each subject, we extracted the average timeseries from each of 419 regions and computed the pair-wise product-moment correlation, resulting in a 419 x 419 functional connectivity matrix, with the upper-triangle of this matrix consisting of 87,571 unique connectivity estimates.

## Statistical analysis

### Characterizing the effects of parenthood and aging on brain function

To characterize the effect of parity (number of children) on brain function, we examined the Spearman correlation between functional connectivity and parity at each edge, using the Network Based Statistic (NBS) for inference at the level of connected components of edges showing a common effect, with significance assessed at *p*_*FWE*_ < 0.05. Spearman correlation was selected for these analyses as both the brain^12,29,30^ and behavioral^101^ effect of parity has shown to be monotonic, rather than strictly linear.

The NBS procedure results in a substantial boost in statistical power compared to mass univariate analysis^102^. The process involves setting a primary component-forming threshold, *τ*, to both the observed and permuted data. Here, *τ* was set to *p* < .01.The choice of this threshold is arbitrary; more lenient thresholds will be sensitive to weaker differences distributed over a large number of edges while more stringent thresholds will be sensitive to stronger effects possibly extending over smaller subsets of edges. One thousand null permutations were computed by randomly resampling parity values across subject, without replacement. For both the observed and permuted null data, the size (number of edges) of the connected components in the supra-thresholded network was recorded. The size of largest component from each permutation was used to build a null distribution of the maximal statistic, which can be used to obtain a FWE-corrected *p*-value for each observed component, as the proportion of null component sizes larger than the observed value. Models were run separately for males and females. Functional connectivity values were adjusted for age, education and TDI, prior to being entered into the model. To ensure the robustness of our findings, we repeated our primary analysis after excluding those with 0 children and without including model covariates. To characterize the effect of age on brain function, we repeated the above procedure using age instead of parity, with models adjusted for education and TDI.

To comprehensively delineate changes in functional connectivity across the 87,571 different connections, we present the results for both age and parity at three different scales: (1) the individual edge level (e.g., Fig1A); (2) the network-level, in which different regions are aggregated into one of eight canonical brain networks^42,57,103^ where we show the proportions of affected edges both within and between these networks (e.g., Fig2A); and, (3) the level of individual brain regions, to identify specific brain areas attached to a high number of connections implicated in the detected NBS network (Fig3).

Different brain networks have intrinsic differences in size, meaning that larger networks will generally have a higher likelihood of being implicated. To determine whether the observed functional connectivity associations showed any network-specificity, we present the proportion of edges within a given NBS component that fell within each of eight brain networks normalized by the total number of possible network connections, which accounts for differences in the number of potential connections (i.e., the tendency for networks with more regions to be more likely to be implicated in a brain-wide analysis).

### Sex consistency in parenthood and aging related brain function

To examine association between males and females on both parity-related and age-related functional connectivity associations, we computed the product-moment correlation between the upper-triangles of the effect-size matrices. For instance, for parity, the unthresholded effect size matrix of spearman *ρ* values between functional connectivity and parity, computed as part of the procedure described above, was vectorized and the upper triangles correlated between males and females. To assess the significance of this association, we used a bootstrapping procedure^58,104^ to build a null distribution of product-moment correlations by repeatedly computing the associations between the observed and null effect using 1000 bootstrapped matrices which were generated by resampling individuals within each sex with replacement and computing associations between brain function and parity across all connections. We also investigated these associations at the network-level, where the observed effect size matrix for each sex were Fishers Z-transformed then averaged within and between 8 network blocks, resulting in 36 network-level effect. The same bootstrapping procedure was followed for network-level effects, where the null effect size matrices were also averaged into network blocks prior to building a null distribution of network-level product-moment correlations. To examine between-sex consistency in ageing effects, the same procedure was repeated for age, instead of parity.

### Association between parenthood and aging related brain function

To examine whether the effect of parity on brain function was in the opposite or same direction as the effect of aging on brain function, for each sex, we computed the product-moment correlation between the ageing and parity effect size matrices across all connections. To assess the statistical significance of this associations, we again used a bootstrapping procedure^58,104^ to build a null distribution of product-moment correlations by repeatedly computing the associations between the observed and null effect using 1000 bootstrapped matrices which were generated by resampling individuals within each sex with replacement and computing associations between brain function and parity and brain function and age across all connections. For each sex, we also examined the association between parity and age-related effects on brain function at the network level by averaging the observed effects within and between network blocks, following the same procedure described above, and that used to examine sex differences in parenthood and aging related brain function.

## Acknowledgements

ERO is supported by a Kavli Institute for Neuroscience Postdoctoral Fellowship, and an American Association for University Women (AAUW) International Fellowship. SC is supported by an American Australian Association Graduate Research Fund Scholarship. This work was supported by the National Institute of Mental Health (Grant Nos. R01MH120080 [to AJH and BTTY] and R01MH123245 [to AJH]). SDJ is supported by a National Health and Medical Research Council of Australia Fellowship APP1174146. HJVR is supported by NIH R01 HD108218, R01 DA050636, and R21 DA052620. BTTY is supported by the NUS Yong Loo Lin School of Medicine (NUHSRO/2020/124/TMR/LOA), the Singapore National Medical Research Council (NMRC) LCG (OFLCG19May-0035), NMRC CTG-IIT (CTGIIT23jan-0001), NMRC STaR (STaR20nov-0003), Singapore Ministry of Health (MOH) Centre Grant (CG21APR1009), the Temasek Foundation (TF2223-IMH-01), and the United States National Institutes of Health (R01MH120080 & R01MH133334). Any opinions, findings and conclusions or recommendations expressed in this material are those of the authors and do not reflect the views of the Singapore NMRC, MOH or Temasek Foundation.

## Supplement

**SFig1.**
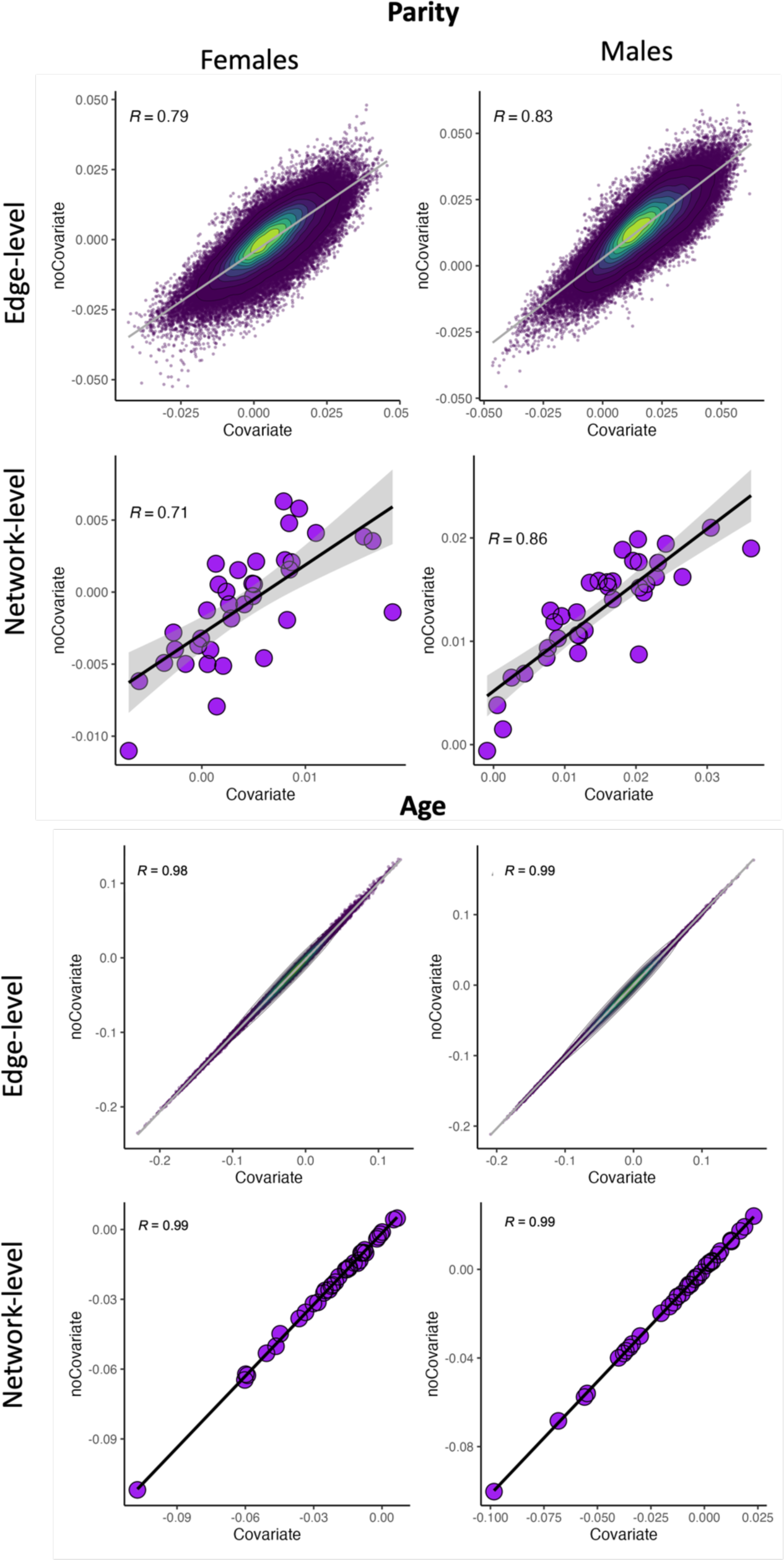
– Comparisons at the edge and network levels between models including covariates vs models not including covariates, for both age and parity.

**SFig2.**
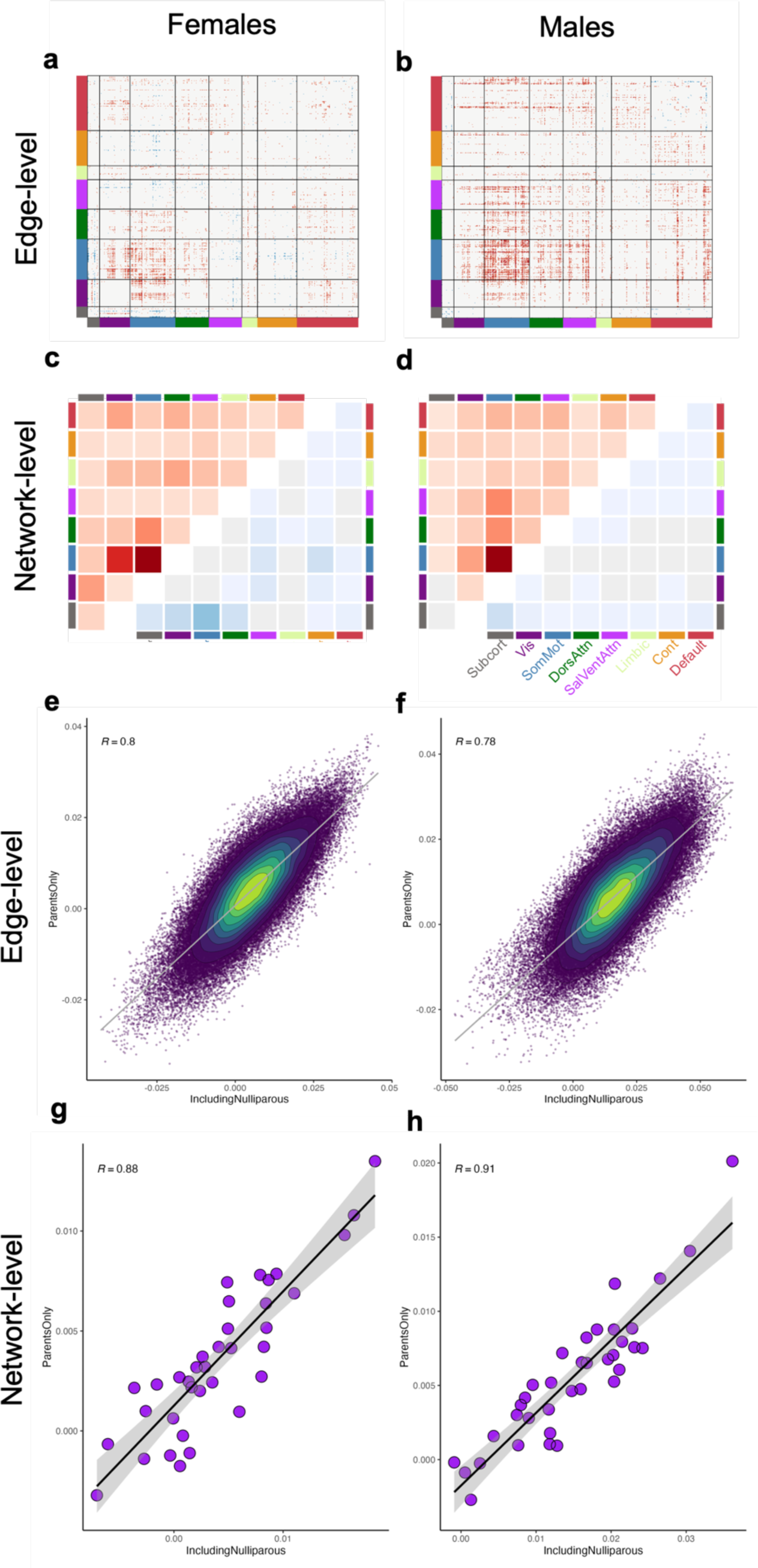
– Comparisons at the edge and network levels between models including those with 0 children vs models not including those with 0 children, for both age and parity.

## Notes

### Competing Interest Statement

The authors have declared no competing interest.

